# SquiggleKit: A toolkit for manipulating nanopore signal data

**DOI:** 10.1101/549741

**Authors:** James M. Ferguson, Martin A. Smith

## Abstract

The management of raw nanopore sequencing data poses a challenge that must be overcome to accelerate the development of new bioinformatics algorithms predicated on signal analysis. SquiggleKit is a toolkit for manipulating and interrogating nanopore data that simplifies file handling, data extraction, visualisation, and signal processing. Its modular tools can be used to reduce file numbers and memory footprint, identify poly-A tails, target barcodes, adapters, and find nucleotide sequence motifs in raw nanopore signal, amongst other applications. SquiggleKit serves as a bioinformatics portal into signal space, for novice and experienced users alike. It is comprehensively documented, simple to use, cross-platform compatible and freely available from (https://github.com/Psy-Fer/SquiggleKit).

## Introduction

Nanopore sequencers, such as those manufactured by Oxford Nanopore Technologies (ONT), generate long sequencing reads by measuring disruptions in ionic current as native biopolymers transit through a nanopore (Garalde *et al.*, 2018; Laszlo *et al.*, 2014). The resulting signal is then de-convoluted into nucleotide sequence through probabilistic models, which often introduce errors given sampling stochasticity and imperfect models (Schreiber and Karplus, 2015; Rang *et al.*, 2018).

When a strand of nucleotides is sequenced with ONT devices, a data file is generated containing the raw signal and machine metadata. Sequencing runs can produce upwards of 100 million reads (depending on the device and sample), which are grouped into a multitude of binary hierarchical data format (.fast5) files. As the yield of ONT sequencers is steadily increasing, discarding raw data once converted into bases is a tempting storage solution. However, until base calling algorithms are developed that can integrate the multitude of known nucleotide analogues (Jonkhout *et al.*, 2017), raw nanopore signal data can be revisited and mined to extract such information (Simpson *et al.*, 2017; Rand *et al.*, 2017; Simpson *et al.*, 2016).

In addition to the need to process a vast quantity of files, extracting pertinent information from the binary file format can be a significant hurdle to overcome when analyzing nanopore data. Once files are processed and data is extracted, nanopore signal data remains difficult to navigate given the the unfamiliar and noisy nature of single molecule sensing. Altogether, these challenges hinder the development of novel bioinformatics tools and training of more accurate machine learning algorithms.

Here we describe SquiggleKit, a toolkit for managing the extensive number of data files, extracting signal data, plotting, and examples of processing the noisy raw signal data. SquiggleKit also acts as a simple guide and starting point for developing new tools which utilise nanopore signal data.

## The Toolkit

SquiggleKit is composed of the follow tools:

### File management and processing

– **Fast5_fetcher** extracts individual fast5 files from an index based on a list of reads of interest, thus reducing both search time and storage space.
– **SquigglePull** opens fast5 files, extracts the embedded signal data, and converts it into a tab separated (.tsv) format.

### Visualisation

– **SquigglePlot** is a command line visualisation tool for signal data.

### Targeting regions of interest in raw signal data

– **Segmenter** identifies the boundaries of relatively long regions of signal attenuation, such as adapter stalls and homopolymer stretches, by measuring the difference in average signal across a sliding window, with error tolerance for signal noise.
– **MotifSeq** identifies raw signal traces that correspond to a given nucleotide sequence, such as an adapter, barcode, or motif of interest. *MotifSeq* takes a query nucleotide sequence as input, converts it to a normalised signal trace (i.e. ‘events’) using Scrappie (https://github.com/nanoporetech/scrappie), then performs signal-level local alignment using a dynamic programming algorithm. *MotifSeq* outputs the location of a matching target in the raw signal with associated metrics.

## Practical example

SquiggleKit can be used to facilitate data management, to generate fine-tuned datasets for machine learning, to visualise signal, to validate demultiplexing results, or to identify motifs of interest without base calling, amongst other applications.

In the following example, we demonstrate how SquiggleKit can be used to validate the 3’ end of a terminal exon using raw nanopore signal data from a cDNA run. Specifically, a cDNA read that aligned to 2 distinct isoforms from a reference transcriptome (**Figure 1A**). We will interrogate the raw signal to identify the 3’ end of one of the reference isoforms.

**Figure 1.**
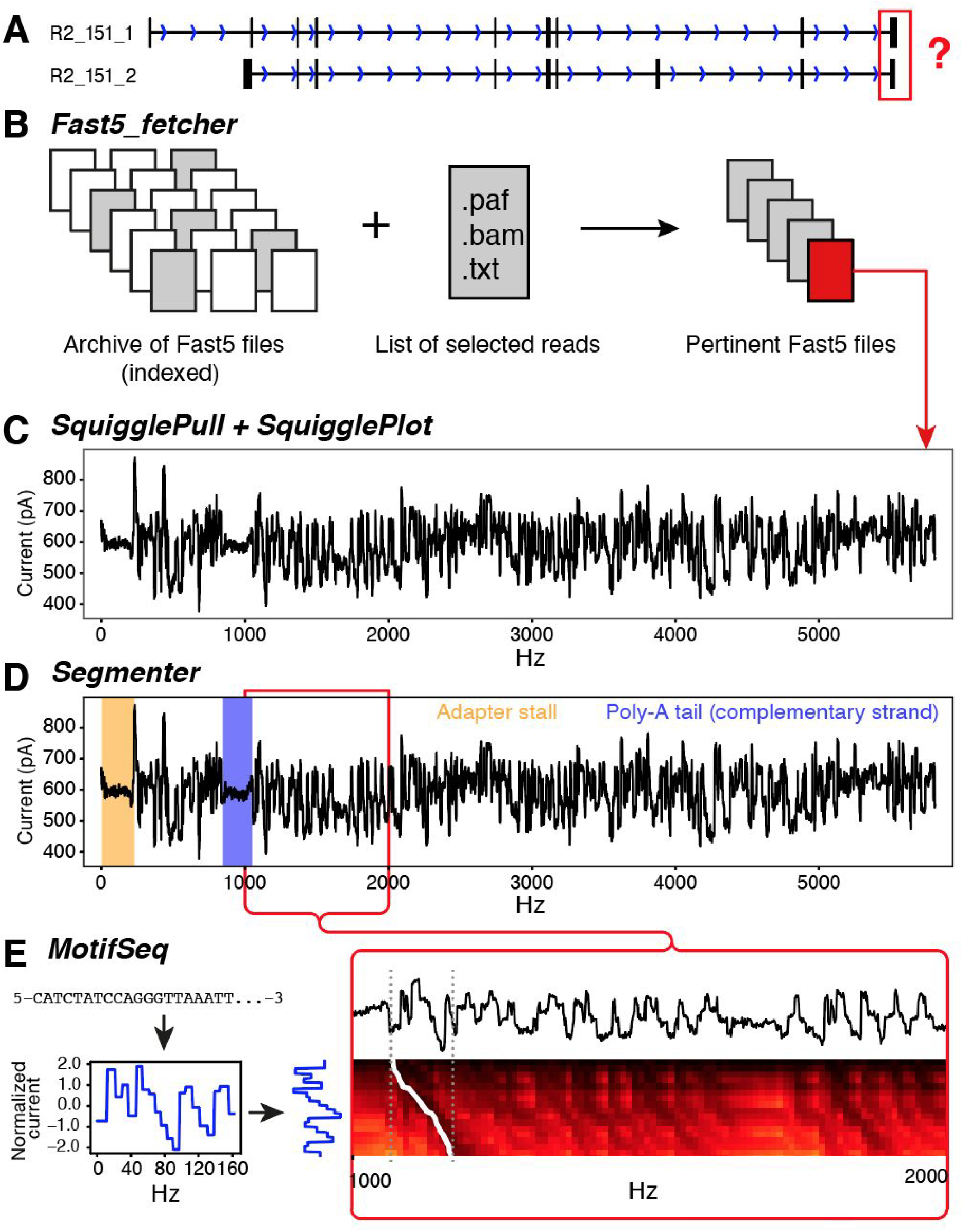
Example usage of SquiggleKit. (**A**) Two similar reference transcripts with similar mapping. (**B**) Using *fast5_fetcher* to extract raw files; (**C**) *SquigglePull* and *SquigglePlot* converting to.tsv and plotting the signal; (**D**) adapter stalls and poly-A tail identified using *Segmenter*; (**E**) converting last 20nt of 3’ end to a signal using a pore current model and aligning this synthetic signal to the empirical signal (dashed grey lines) by backtracking through a dynamic programming matrix (white trace).

First, a list of all read identifiers that mapped to isoform *A* are extracted from the alignment output into a.txt file. The list is used as input for *Fast5_fetcher* (**Figure 1B**) together with an index of all archived raw data files (.fast5.tar), resulting in 45.fast5 files being extracted instead of all 2,710,372 in the full dataset.

Next, the raw ionic signal for the read of interest was extracted using *SquigglePull* and visualised with *SquigglePlot* (**Figure 1C**). The.tsv output of *SquigglePull* is then used as input for *Segmenter* to identify the adapter stall and poly-A homopolymer sequence (**Figure 1D**). The raw signal directly downstream of the the poly-A signal is then extracted with *Segmenter*, thus selecting the 3’ end of the terminal exon in the signal data.

Finally, the nucleotide sequence corresponding to the last 20 nucleotides of the 3’ end of isoform *A* is converted into normalized signal space via *MotifSeq*. The resulting theoretical current trace is aligned to the empirical signal from *Segmenter* (**Figure 1E**), returning the position of the sequence in the terminus of the raw signal, and confirming the boundary of the exon.

## Conclusion

SquiggleKit helps researchers extract more information from their data by making raw signal analysis more accessible. Unlike more specialized tools for nanopore signal visualization and analysis, such as Tombo (Stoiber *et al.*, 2016) and BulkVis (Payne *et al.*, 2018), SquiggleKit is tailored towards broad applications and user adaptability. We anticipate that this set of applications will facilitate the development of future bioinformatics tools and help create more accurate probabilistic models for nanopore sequencing data analysis, in a timely and user friendly manner.

## Acknowledgements

We would like to thank Shaun Carswell, Hasindu Gamaarachchi, Kai Martin, and Tansel Ersavas for testing the toolkit and providing valuable feedback. Many thanks to Tim Mercer who kindly provided the synthetic RNA controls used in the practical example.

## References

Garalde, D.R. et al. (2018) Highly parallel direct RNA sequencing on an array of nanopores. Nat. Methods, 15, 201–206.

Jonkhout, N. et al. (2017) The RNA modification landscape in human disease. RNA, 23, 1754–1769.

Laszlo, A.H. et al. (2014) Decoding long nanopore sequencing reads of natural DNA. Nat. Biotechnol., 32, 829–833.

Payne, A. et al. (2018) BulkVis: a graphical viewer for Oxford nanopore bulk FAST5 files. Bioinformatics.

Rand, A.C. et al. (2017) Mapping DNA methylation with high-throughput nanopore sequencing. Nat. Methods, 14, 411–413.

Rang, F.J. et al. (2018) From squiggle to basepair: computational approaches for improving nanopore sequencing read accuracy. Genome Biol., 19, 90.

Schreiber, J. and Karplus, K. (2015) Analysis of nanopore data using hidden Markov models. Bioinformatics, 31, 1897–1903.

Simpson, J.T. et al. (2017) Detecting DNA cytosine methylation using nanopore sequencing. Nat. Methods, 14, 407.

Stoiber, M.H. et al. (2016) De novo Identification of DNA Modifications Enabled by Genome-Guided Nanopore Signal Processing. BioRxiv doi.org/10.1101/094672

